# Computationally defined and *in vitro* validated putative genomic safe harbour loci for transgene expression in human cells

**DOI:** 10.1101/2021.12.07.471422

**Authors:** Matias I. Autio, Efthymios Motakis, Arnaud Perrin, Talal Bin Amin, Zenia Tiang, Dang Vinh Do, Jiaxu Wang, Joanna Tan, Wei Xuan Tan, Adrian Kee Keong Teo, Roger S.-Y. Foo

## Abstract

Selection of the target site is an inherent question for any project aiming for directed transgene integration. Genomic safe harbour (GSH) loci have been proposed as safe sites in the human genome for transgene integration. Although several sites have been characterised for transgene integration in the literature, most of these do not meet criteria set out for a GSH and the limited set that do have not been characterised extensively. Here, we conducted a computational analysis using publicly available data to identify 25 unique putative GSH loci that reside in active chromosomal compartments. We validated stable transgene expression and minimal disruption of the native transcriptome in three GSH sites *in vitro* using human embryonic stem cells (hESCs) and their differentiated progeny. Furthermore, for easy targeted transgene expression, we have engineered constitutive landing pad expression constructs into the three validated GSH in hESCs.

## Introduction

Stable expression of transgenes is essential in both therapeutic and research applications. Traditionally, transgene integration has been accomplished via viral vectors in a semi-random fashion, but with inherent integration site biases linked to the type of virus used (Mitchell et al., 2004). The randomly integrated transgenes may undergo silencing (Ellis, 2005; Mok et al., 2007) and more concerningly, can also lead to dysregulation of endogenous genes. Gene dysregulation can lead to malignant transformation of cells and has unfortunately given rise to cases of leukaemia (Hacein-Bey-Abina et al., 2003; Howe et al., 2008) in gene therapy trials. GSH loci have been previously suggested as safe sites for transgene integration. Criteria proposed for a putative GSH include; a set distance from coding and non-coding genes; with added separation from known oncogenes and miRNAs, and no disruption of transcriptional units or ultraconserved regions (Papapetrou and Schambach, 2016; Papapetrou et al., 2011; Sadelain et al., 2012). To date, a number of sites in the human genome have been used for directed integration; however none of these pass scrutiny as *bona fide* GSH (Papapetrou and Schambach, 2016). Here, we conducted a computational analysis to filter sites that meet criteria for GSH loci. In addition to the safety criteria, we identified regions that reside in active chromosomal compartments in many human cell and tissue types. Our analysis yielded a final list of 25 unique putative GSH that are predicted to be accessible in multiple cell types. We used human embryonic stem cells (hESCs) and their differentiated progeny to validate stable transgene expression in three of the putative GSH sites *in vitro*. Furthermore, to enable easy targeted transgene expression, we generated three hESC lines with constitutive landing pad expression constructs targeted into the three validated GSH.

## Results

### Computational filtering for safe and accessible loci

A list of criteria has previously been suggested for a given locus to qualify as putative GSH (Papapetrou and Schambach, 2016; Sadelain et al., 2012). These criteria state that GSH: must not be in proximity to genes coding or non-coding, with added distance from known oncogenes and miRNAs, and must not disrupt transcriptional units or ultra-conserved genomic regions. To shortlist putative GSH we conducted a computational search of the human genome using publicly available data (Fig. 1 A). We included the previously published safety criteria and added a further filter to exclude any regions of DNaseI hypersensitivity, as these regions are likely enriched in transcription factor binding and regulatory elements (Meuleman et al., 2020). A total of 12,766 sites, ranging from 1 b to approximately 30 Mb, passed the filters used (Fig. 1 A-B, Supp. Table 1). For a universal GSH site to be useful, it needs not only to be safe, but also enable stable expression of a transgene in any tissue type. We filtered the human genome for regions consistently in the active chromatin compartment based on 21 different human cell and tissue types (Schmitt et al., 2016). To extend the analysis beyond the limited set of samples, we utilised RNA-seq data of all available tissue types from the GTEx portal (Carithers et al., 2015). We selected an empirical set of ubiquitously expressed genes with low variance. We then cross referenced the chromosomal locations of these genes with the consistently active chromatin regions. This analysis yielded 399 1Mb active regions that overlapped with a ubiquitously expressed gene (Fig. 1 A-B, Supp. Table 1). By overlapping the two datasets, we found 49 safe sites within the active regions. We further filtered the 49 sites (Fig. 1 A-B, Supp. Table 1) using BLAT to generate a final shortlist of 25 unique putative universal GSH sites in the human genome (Fig. 1 A-B, Table 1).

**Table 1:** Coordinates of candidate GSH, their associated active chromosome regions & housekeeping gene, and BLAT score against the most similar region

| Chromosome | Start | End | Width | Active region start | Active region end | Housekeeping gene | BLAT-score |
| --- | --- | --- | --- | --- | --- | --- | --- |
| 1 | 113289036 | 113289342 | 307 | 113000001 | 114000000 | HIPK1 | 0.21 |
| 1 | 113314841 | 113318369 | 3529 | 113000001 | 114000000 | HIPK1 | 0.18 |
| 1 * | 113339961 | 113340514 | 554 | 113000001 | 114000000 | HIPK1 | 0.27 |
| 2 * | 128912721 | 128914814 | 2094 | 128000001 | 129000000 | UGGT1 | 0.08 |
| 2 | 128918961 | 128919839 | 879 | 128000001 | 129000000 | UGGT1 | 0.16 |
| 2 * | 128932307 | 128935799 | 3493 | 128000001 | 129000000 | UGGT1 | 0.15 |
| 2 | 128963272 | 128965759 | 2488 | 128000001 | 129000000 | UGGT1 | 0.44 |
| 2 | 208992998 | 208997459 | 4462 | 208000001 | 209000000 | PIKFYVE | 0.28 |
| 4 * | 17373361 | 17374159 | 799 | 17000001 | 18000000 | MED28 | 0.04 |
| 5 | 131058585 | 131058947 | 363 | 131000001 | 132000000 | FNIP1 | 0.41 |
| 5 | 148753741 | 148757219 | 3479 | 148000001 | 149000000 | FBXO38 | 0.10 |
| 6 * | 15727241 | 15727490 | 250 | 15000001 | 16000000 | DTNBP1 | 0.47 |
| 7 | 4314741 | 4315279 | 539 | 4000001 | 5000000 | FOXK1 | 0.15 |
| 7 | 4321017 | 4323839 | 2823 | 4000001 | 5000000 | FOXK1 | 0.09 |
| 7 | 4328040 | 4329659 | 1620 | 4000001 | 5000000 | FOXK1 | 0.21 |
| 7 | 4353504 | 4354219 | 716 | 4000001 | 5000000 | FOXK1 | 0.32 |
| 7 | 4454808 | 4456201 | 1394 | 4000001 | 5000000 | FOXK1 | 0.17 |
| 8 | 23945241 | 23945819 | 579 | 23000001 | 24000000 | R3HCC1 | 0.34 |
| 8 | 23986981 | 23988319 | 1339 | 23000001 | 24000000 | R3HCC1 | 0.06 |
| 8 | 23999628 | 24001194 | 1567 | 23000001 | 24000000 | R3HCC1 | 0.02 |
| 18 | 56339813 | 56340245 | 433 | 56000001 | 57000000 | TXNL1 | 0.06 |
| 18 | 56396821 | 56397319 | 499 | 56000001 | 57000000 | TXNL1 | 0.07 |
| 18 | 56410681 | 56411039 | 359 | 56000001 | 57000000 | TXNL1 | 0.14 |
| 18 * | 56534775 | 56536439 | 1665 | 56000001 | 57000000 | TXNL1 | 0.16 |
| 19 * | 5400761 | 5402139 | 1379 | 5000001 | 6000000 | SAFB | 0.18 |

\* = GSH shortlisted for *in vitro* validation

**Figure 1.**
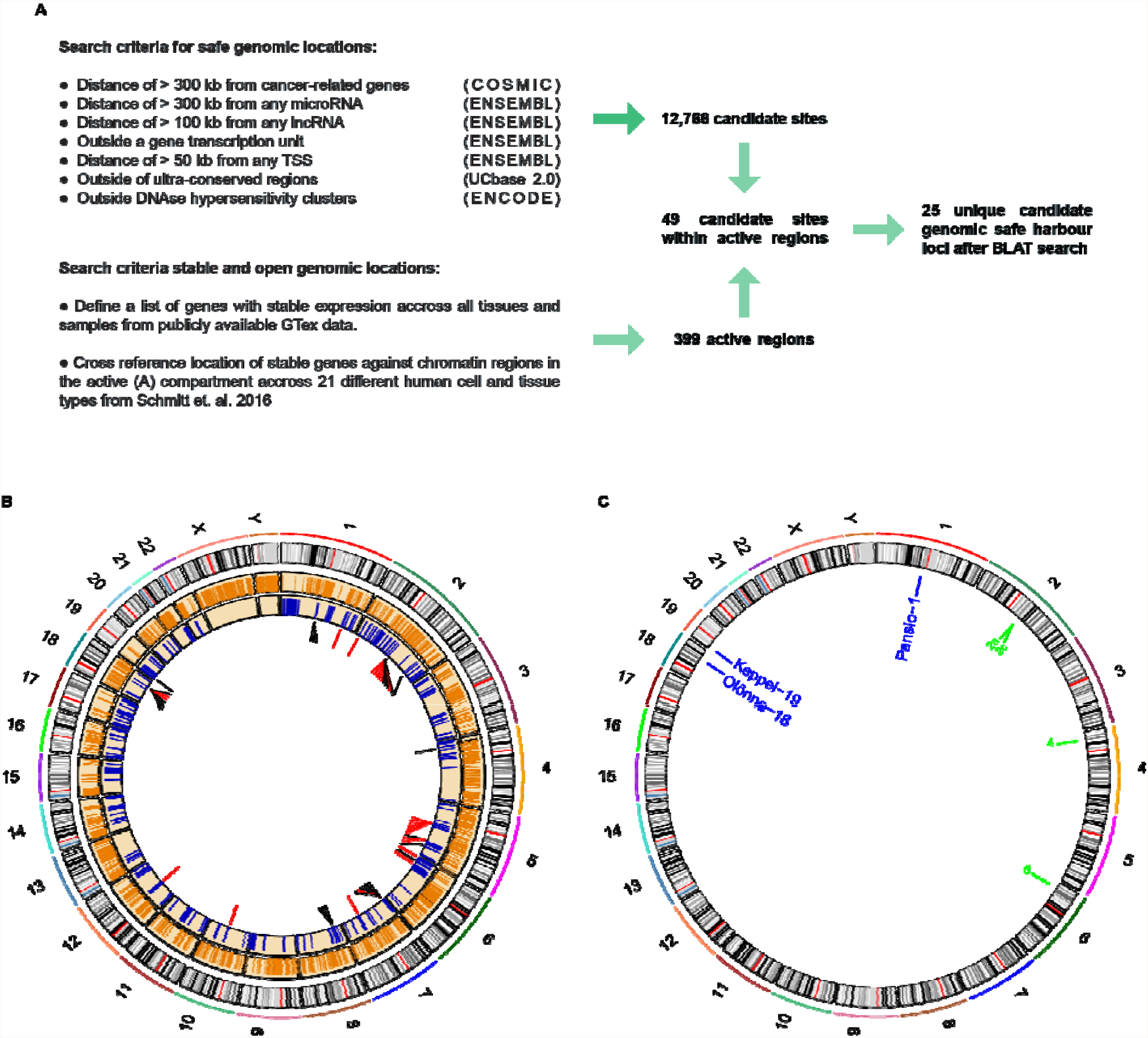
**A)** Schematic representation of the computational workflow for defining candidate GSH. **B)** CIRCOS plot summarising computational search results. Ring 1: chromosome ideograms; ring 2: orange bars indicating safe sites; ring 3: blue bars indicating active regions; ring 4: candidate sites within active regions, red bars site failed BLAT screening, black bars site passed BLAT screening. **C**) Locations of candidate GSH targeted *in vitro*. Blue labels: targeted clone established; green labels: no clone established.

### Targeted knock-in at putative GSH with CRISPR/Cas9

To validate our candidate GSH, we selected seven of the 25 sites for *in vitro* experiments (Fig. 1 C). None of the selected seven sites lie at or immediately adjacent to borders of topologically associated domains (TADs) (Supp. Fig. 1). We targeted H1 hESC using CRISPR/Cas9 and a donor landing pad construct (Fig. 2 A) at each of the seven candidate sites (Methods and Supp. Table 2). To exclude potential off-target effects of CRISPR/Cas9 targeting, we used a version of Cas9 with enhanced specificity (Slaymaker et al., 2016) and effective guides with highest predicted specificity available. Following antibiotic selection, single clones were expanded and screened for successful homology directed repair driven integration of the expression construct with junction- and digital-PCR (Fig. 2 B, Supp. Table 3, and Supp. Fig. 2). Successful heterozygous targeting of the donor construct was confirmed at three candidate GSH sites on chromosomes 1, 18 and 19. No evidence of off-target activity was observed following PCR amplification and Sanger sequencing of the top 5 predicted off-target sites for each of the targeted clones (Supp. Fig. 3). We named the successfully targeted safe harbours after real world harbours, designating them Pansio-1, Olônne-18, and Keppel-19.

**Figure 2.**
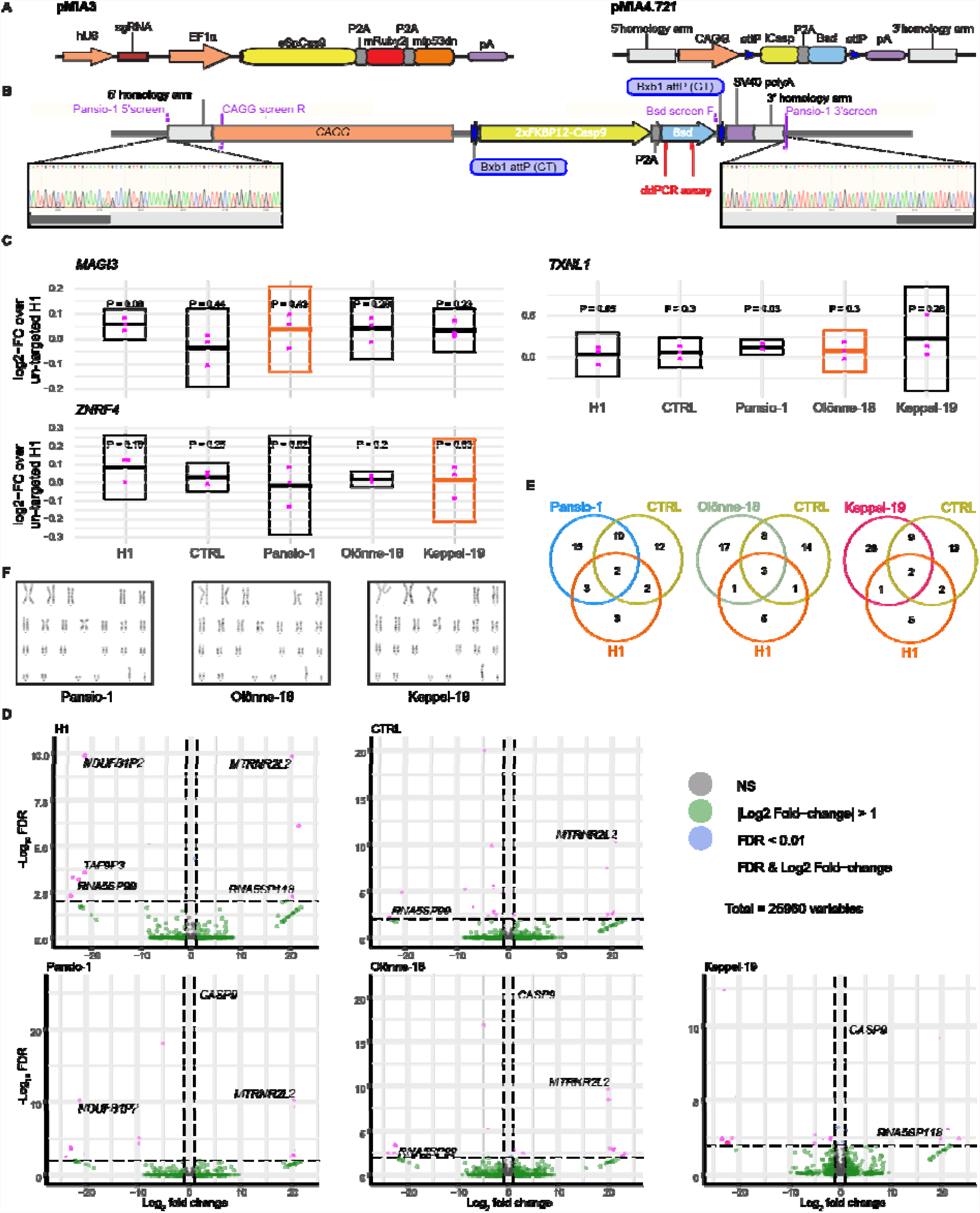
**A)** Schematic representation of CRISPR/Cas9 plasmid (pMIA3) and homology directed repair donor (pMIA4.721) used for targeting with functional components annotated. **B)** Schematic of integrated landing pad expression construct. Positions of primers for junction-PCR as well as of ddPCR assay are indicated. Representative junction-PCR Sanger sequencing reads from Pansio-1 targeted clones shown in expanded view. **C**) Log2-FC of mRNA expression levels against un-targeted H1 hESC samples for the nearest genes of Pansio-1, Olônne-18, and Keppel-19 candidate GSH. Evaluated samples: H1 = un-targeted hESC, CTRL = H1 cells non-GSH targeted, Pansio-1 = landing pad construct integrated to Pansio-1 GSH in H1 hESC, Olônne-18 = landing pad construct integrated to Olônne-18 GSH in H1 hESC and Keppel-19 = landing pad construct integrated to Keppel-19 GSH in H1 hESC. Box plots representing 95% confidence intervals of mean log2-FC. Nearest gene for each GSH indicated in orange. Individual data points shown in pink with P-value for each comparison shown above. **D)** Volcano plots of RNA-seq analysis against un-targeted H1 hESC. Samples analysed as in **C)**. Differentially expressed (DE) genes with FDR ≤ 0.01 and |logFC| ≥ 1 in pink, genes with |logFC| ≥ 1 in green, genes with FDR ≤ 0.01 in blue, others in grey. E) Venn-diagrams illustrating the overlap of DE genes between un-targeted H1 hESC, non-GSH targeted H1 hESC and the three GSH targeted H1 hESC lines. **F**) Representative images of metaphase spreads used for karyotyping the GSH targeted cell lines.

### *In vitro* validation of targeted GSH in hESCs

To investigate the safety of our targeted GSH, we first checked the mRNA expression levels of the nearest genes *MAGI3, TXNL1* and *ZNRF4* to Pansio-1, Olônne-18 and Keppel-19 respectively using qPCR. When compared to un-targeted H1 hESCs, the 95% confidence intervals of log2 fold-change (log2-FC) for each gene in the respective GSH line overlapped with zero, indicating that the data do not show evidence of statistically significant change in mRNA expression levels of the nearest genes (Fig. 2 C). We then conducted RNA-seq analysis to look for gene expression changes on a global scale. Our GSH targeted clones yielded very low numbers of differentially expressed (DE) genes; 30, 29 and 40 respectively for Pansio-1, Olônne-18, and Keppel-19 (Fig. 2 D and Supp. Table 4). Notably *CASP9*, the suicide gene included in our targeting construct, was the gene with lowest false discovery rate (FDR) in Pansio-1 and Olônne-18 and second lowest in Keppel-19 (Fig. 2 D and Supp. Table 4). A high proportion of the DE genes found in our GSH targeted lines were shared with control targeted H1 cells and/or untargeted wild-type H1 cells (Fig. 2 E and Supp. Table 4), suggesting that the observed changes were unrelated to the GSH targeting and transgene expression. The closest DE gene found from our analysis lies >42 Mb away for Pansio-1 and >47 Mb for Keppel-19. For Olônne-18 there were no DE genes on the same chromosome (Supp. Table 4). Functional enrichment analysis of the DE genes revealed relatively few terms; 9, 8 and 15 for Pansio-1, Olônne-18, and Keppel-19 respectively (Supp. Table 4). In the case of Pansio-1 all the terms were shared with the control targeted H1 cells except for one, which was due to the increased level of *CASP9* (Supp. Table 4). As a further safety check, we also conducted karyotyping of the three targeted clones and observed no abnormalities (Fig. 2 F). Taken together, these results suggest minimal disruption of the native genome and no karyotypic abnormalities following transgene integration to our three GSH.

In addition to safety, a functional GSH also needs to allow for stable expression of a transgene. We took advantage of the landing-pad design of our targeting construct to swap in a sequence coding for Clover-fluorophore, by introducing a plasmid expressing BxbI-integrase as well as a donor construct into the three GSH lines (Fig. 3 A-B). Targeted cells were enriched with fluorescence activated cell sorting. Introduction of the payload transgene did not alter the expression of pluripotency markers OCT3/4 and SOX2 (Fig. 3C-D). We maintained the Clover targeted GSH lines in hESC state over 15 passages and consistently observed >98% Clover positive cells (Fig. 3E). Interestingly, the levels of Clover integrated to Olônne-18 seemed to be consistently lower than integrations to Pansio-1 or Keppel-19 (Fig. 3 C-E). To investigate the stability of our GSH in other cell types, we conducted directed differentiation of our Clover-integrated hESC lines into cell types from the three germ lineages. Clover expression remained consistent in neuronal, liver, and cardiac cells (Fig. 3F-H, Supp. Fig. 4) in Pansio-1, Olônne-18, and Keppel-19 targeted cells.

Overall, we have developed a computational pipeline to define GSH candidate sites from the human genome that fulfil criteria for safety as well as accessibility for transgene expression. Our pipeline defines 25 unique candidate GSH and we conducted *in vitro* validation experiments for three of them, Pansio-1, Olônne-18, and Keppel-19. Targeting and transgene expression in hESC at the three sites led to minimal or no change in the expression levels of the nearest native genes or the transcriptome overall and did not interfere with directed differentiation to the three germ lineages. Furthermore, we established landing pad expression lines in H1 hESC of Pansio-1, Olônne-18, and Keppel-19, which we hope will serve as useful research tools.

**Figure 3.**
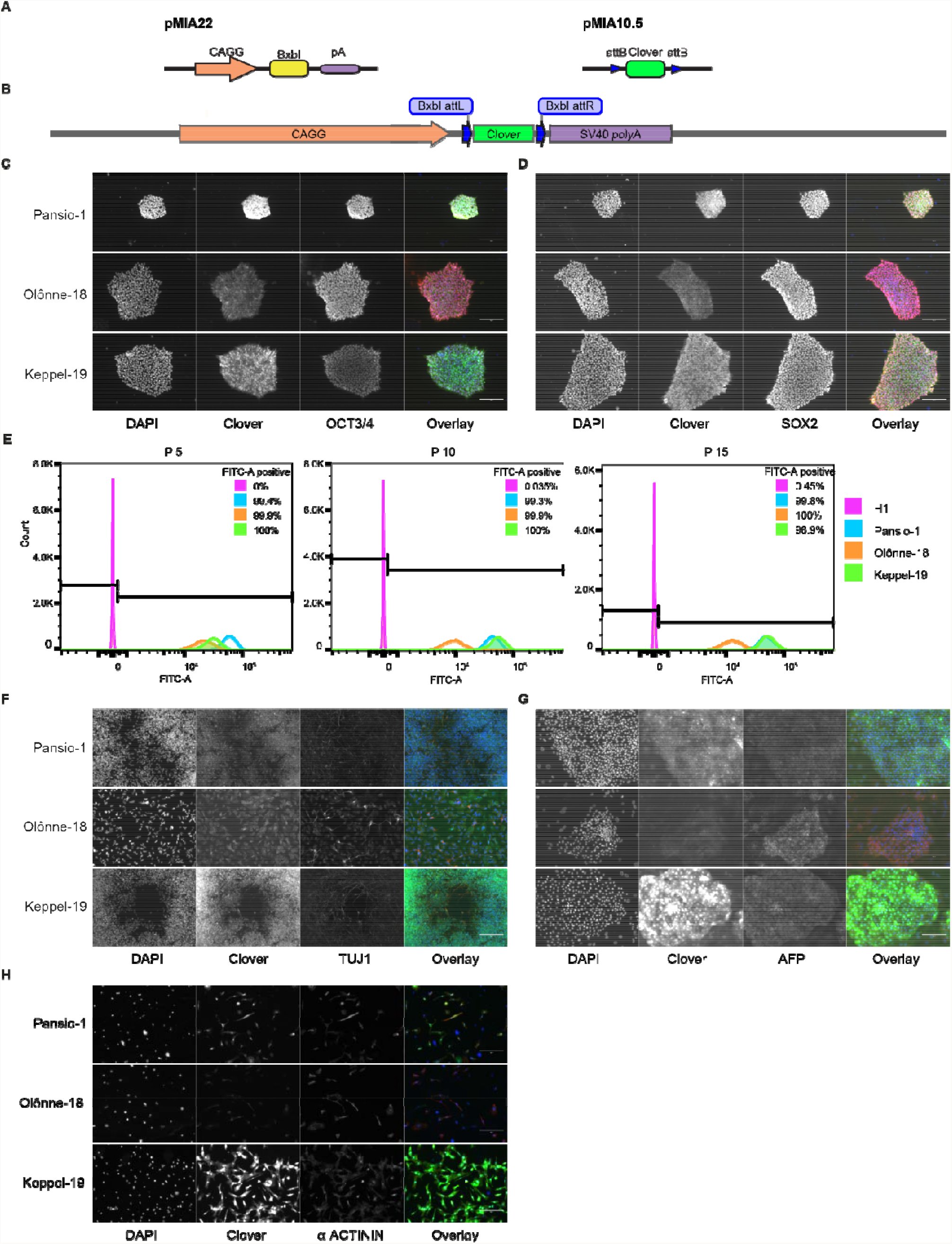
**A)** Schematic representation of integrase expression construct (pMIA22) and transposon donor construct (pMIA10.5). **B)** Schematic of landing pad construct with integrated Clover transgene. **C)** Representative immunofluorescence images of Clover-integrated GSH H1 cells. DAPI = nuclear staining with 4′,6-diamidino-2-phenylindole, Clover = fluorescence from Clover transgene, OCT3/4 = antibody staining against OCT3/4, Overlay = overlay of the three imaged channels. **D)** As in **C)** apart from antibody staining against SOX2. **E)** Histograms of flow cytometry analysis for FITC-A channel of un-targeted H1 hESC, and the three GSH targeted hESC lines over 15 passages. Percentages of FITC-A positive cells according to the indicated gating. **F)** Representative immunofluorescence images of Clover-integrated GSH H1 cells differentiated to neuronal-like cells. Channels imaged as in **C)** apart from antibody staining against Tuj1. **G)** As in **F)** for cells differentiated to hepatocyte-like cells, antibody staining against AFP. **H)** As in **F)** for cells differentiated to cardiomyocyte-like cells, antibody staining against sarcomeric α-ACTININ. Scale bars for all immunofluorescence images equal to 150 μm.

## Discussion

A GSH site is an ideal location for transgene integration. To qualify as a GSH, a locus should be able to host transgenes enabling their stable expression as well as not interfere with the native genome (Papapetrou and Schambach, 2016). A number of previous studies have reported discovery and usage of a handful of integration sites (Costa et al., 2005; Eyquem et al., 2013; Papapetrou et al., 2011; Pellenz et al., 2019; Rodriguez-Fornes et al., 2020), which fulfil a subset of criteria previously suggested for GSH (Papapetrou and Schambach, 2016; Sadelain et al., 2012). However, in contrast to the candidate sites we present (Fig. 1 A-B), the previously reported sites do not utilise criteria to avoid potential regulatory elements or criteria for universally stable and active genomic regions (Aznauryan et al., 2021; Costa et al., 2005; Eyquem et al., 2013; Papapetrou et al., 2011; Pellenz et al., 2019; Rodriguez-Fornes et al., 2020). We utilised directed differentiation of our Pansio-1, Olônne-18 and Keppel-19 targeted hESC to show consistent expression in hESC and cells from all three germ lineages (Fig. 3 C-H), whereas majority of the previously reported sites remain studied in only a limited number of cell types (Aznauryan et al., 2021; Eyquem et al., 2013; Pellenz et al., 2019; Rodriguez-Fornes et al., 2020).

The integration sites that have been most heavily utilised in research, namely AAVS1, CCR5 and Rosa26 orthologous site (Irion et al., 2007; Kotin et al., 1992; Liu et al., 1996; Perez et al., 2008), do not meet the criteria set out for GSH (Papapetrou and Schambach, 2016; Sadelain et al., 2012). These sites reside in highly gene rich regions and in the case of AAVS1 actually within a gene transcription unit, furthermore all of these loci have known oncogenes in their proximity (<300kb) (Sadelain et al., 2012). Thus, their utilisation in a clinical setting would require extensive further safety data. Furthermore variable transgene expression and silencing has been reported for AAVS1 in hepatocytes (Ordovás et al., 2015) and cardiomyocytes (Bhagwan et al., 2019). The three candidate sites we tested, Pansio-1, Olônne-18, and Keppel-19, demonstrated stable expression of the Clover transgene in both hepatocyte- and cardiomyocyte-like cells after differentiation from hESC (Fig. 3 G-H). The transcriptome analysis of our GSH targeted hESC lines showed very low number of differentially expressed genes (Fig. 2 D) when compared to untargeted H1 hESC. Furthermore, many of the DE genes we observed were shared with independent wild type H1 hESC as well as control targeted H1 hESC (Fig. 2 E). The nearest observed DE genes to each targeted candidate GSH were located >42 Mb away from the targeted site, which is far beyond the distance generally suggested for enhancer-promoter interactions and TADs (Jerkovic’ and Cavalli, 2021).

Our data suggest that the three candidate GSH, Pansio-1, Olônne-18, and Keppel-19, are able to support stable transgene expression in different cell types, and integration at these sites shows minimal perturbation of the native genome. As hESC currently offer the best available model to test the stability of transgene expression from a GSH in the human genome, we generated landing-pad H1 hESC lines for all three candidate sites. These cell lines will allow easy integration of various transgenes to the candidate GSH for research applications. Advances in genome engineering technologies using e.g. novel integrases or CRISPR RNA-guided transposons (Anzalone et al., 2021; Durrant et al., 2021; Ioannidi et al., 2021; Klompe et al., 2019) will likely lead to easier integration of more complex transgene constructs. We hope that our list of 25 candidate GSH and the three *in vitro* validated sites will serve as a resource for research applications in many different cell types. After further experimentation and reproducibility validations, translation to clinical applications can be envisaged.

