## Supplementary material for "Computationally defined and *in vitro* validated putative genomic safe harbour loci for transgene expression in human cells": methods

**STAR Methods**

**Short-listing of putative safe harbour genomic regions**

We applied a series of computational, whole-genome loci filtering criteria to pick a narrow list of high-confidence, putative safe harbour sites for experimental validation. In the first step, we selected genomic regions that satisfy simultaneously all of the following criteria: the loci should be located outside of ultra-conserved regions (Lomonaco et al., 2014; Taccioli et al., 2009) (coordinates lifted over from hg19 to hg38 assembly), outside of DNase clusters +/- 2kb (ENCFF503GCK, ENCODE database <https://www.encodeproject.org/> (Meuleman et al., 2020)), more than 50 kb away from any transcription start site and outside a gene transcription unit (ENSEMBL Release 103, dataset *hsapiens_gene_ensembl*, <http://www.ensembl.org/index.html>), more than 300 kb away from cancer-related genes (Cancer Gene Census, GRCh38, COSMIC v92 database <https://cancer.sanger.ac.uk/census>), more than 300 kb away from any miRNA (ENSEMBL Release 103, dataset *hsapiens_gene_ensembl*, gene_biotype = miRNA, <http://www.ensembl.org/index.html>) and more than 100 kb away from any long non-coding RNA (ENSEMBL Release 103, dataset *hsapiens_gene_ensembl*, gene_biotype = lncRNA, <http://www.ensembl.org/index.html>). From the filtered loci we discarded the loci with high BLAT similarity to other sequences. On the RNA-seq level, we required the putative safe harbour sites to be associated with ubiquitously expressed, low variance genes. On the 3D chromosome organization level, they should belong to regions consistently located in active chromosomal compartments across multiple tissue types.

**Ubiquitously expressed and low-variance genes**

We downloaded the median gene-level TPMs by tissue type from GTEx (<https://www.gtexportal.org/home/datasets>) and identified an empirical set of low-variance housekeeping genes. To this extent, we estimated the mean and the variance of each gene across all available tissue types and, independently, selected the genes with the lowest, insignificant variability using the HVG function of scran R package that decomposes the total variance of each gene into its biological and technical components. We picked the genes whose expression levels do not change significantly across the tissue types ($FDR>0.9$) and fit a mean vs variance non-parametric lowess regression model. We selected the genes with mean TPM $\geq5$ and variance below the average (smoothed) variance estimated from the lowess model.

**Loci interaction via chromatin conformation capture**

Using a set of publicly available Hi-C chromatin organization data (Schmitt et al., 2016) from human cell and tissue types, we shortlisted the genomic regions consistently located (at least 20/21 interrogated tissue types) in active (open chromatin) compartments (Lieberman-Aiden et al., 2009).

**BLAT analysis**

We measured the uniqueness of target loci using BLAT (<https://genome.ucsc.edu/cgi-bin/hgBlat>) on the human genome GRCh38 with BLAT’S guess query type. BLAT takes the target DNA sequence as input and identifies similar ones in the whole human genome. Target sequences of more than 25,000 bps (BLAT’s limit) were split into multiple smaller overlapping segments of length between 9,000 to 11,000 bps each (depending on the original target length) and tested separately. We reported the location and length of the matching sequences, their similarity to the target (BLAT score) and the number of the matching base pairs. We filtered out the target loci with at least one matching sequence of more than 50% similarity.

**TAD boundary check**

We checked the locations of our *in vitro* targeted GSH candidates against the TAD borders using data from H1 human embryonic stem cells (hESCs) (Dixon et al., 2015) on the 3D Genome Browser (<http://3dgenome.org>) (Wang et al., 2018). Visual inspection of the candidate loci confirmed that all the candidate GSH are more than 80,000 bp away from TAD borders (Supp. Fig. 1).

**Plasmid construction**

All restriction enzymes were purchased from NEB. PCR reactions were conducted using Q5® Hot Start High-Fidelity 2X Master Mix (NEB, M0494L). Ligations were conducted using isothermal assembly with NEBuilder® HiFi DNA Assembly Master Mix (NEB, E2621L). All gBlocks, primers & oligos were ordered from Integrated DNA Technologies, Singapore. Plasmids used in this manuscript will be made available via Addgene. Primers used for fragment amplification are listed in Supp. Table 2.

**pMIA4.721:** An in-house expression plasmid containing a CAGG promoter (pMIA4.9) was digested with BamHI & SphI. Two gBlocks (bxb-bsd and bxb-sv) were directly ligated into the digested plasmid. The resulting plasmid was digested with PmlI & KpnI. A fragment containing codon optimised iCasp9-2A-Bsd was amplified from a plasmid supplied by Genewiz (sequence of iCasp9 based on Straathof *et.al.* (2005) (Straathof et al., 2005)) and ligated into the digested plasmid. This plasmid was subsequently digested with AgeI & SbfI. The SV40 polyA signal was amplified from pMAX-GFP and ligated to the digested plasmid to generate pMIA4.271.

To generate the HDR donors for each GSH candidate, the pMIA4.721 plasmid was digested with NheI for 5’ homology arm and with SbfI for 3’ homology arm. Homology arms ranging from 240bp to 769bp were amplified from H1 hESC gDNA. Ligation of homology arms was done in two sequential reactions. Order of ligation depended on the underlying sequence of homology arms for each target. Plasmid for control targeting was built by ligating 5’ homology arm to pMIA4.9 digested by NheI and 3’ by SbfI.

**pMIA22:** pMAX-GFP (Lonza) was digested with KpnI and SacI. A gBlock encoding a codon optimised BxbI-integrase (Ghosh et al., 2005) with a C-terminal bi-partite nuclear localisation signal(Wu et al., 2009) was amplified and ligated to the digested backbone.

**pMIA10.5-Clover:** An empty donor plasmid (pMIA10.5) containing a 5’ BxbI attB (CT) and a 3’BxbI attB (GT) was ordered from Genewiz. Clover transgene was amplified and ligated into the plasmid after digestion with AgeI & KpnI.

**Stem cell culture**

Human ESC line H1 was maintained using mTeSR medium (Stemcell Technologies, 85850) on 1:200 Geltrex (Thermo fisher, A1413202) coated tissue culture plates and passaged regularly as cell aggregates every 4-5 days using ReLeSR (Stemcell Technologies, 05872).

**CRISPR/Cas9 mediated targeted construct integration in hESC**

H1 hESCs were targeted via nucleofection using an Amaxa-4D (Lonza) as described previously (Ang et al., 2018). Briefly, gRNAs were designed using CRISPOR (Concordet and Haeussler, 2018) (<http://crispor.tefor.net/>). Three gRNAs with the highest predicted off-target scores and containing a native G-base in the first position were selected for each GSH candidate (Supp. Table 2). The gRNAs were cloned into pMIA3 plasmid (Addgene #109399) digested with Esp3I and tested via a GFP reconstitution assay in HEK239T cells. The target locus of each candidate GSH was amplified from H1 hESC gDNA (for primers see Supp. Table 2). The amplified target sequences, ranging from 232-974bp were cloned into the pCAG-EGxxFP plasmid (Mashiko et al., 2013) (Addgene # 50716, a kind gift from Dr Masahito Ikawa). The pMIA3 with the tested gRNA and the respective pCAG-EGxxFP target plasmid were transfected into HEK293T using lipofectamine3000 (ThermoFisher Scientific, L3000015), according to manufacturer’s recommendations. For each candidate GSH the gRNA with the highest GFP signal at 48h post transfection (data not shown) was selected for use in hESC targeting.

Five micrograms of pMIA3 plasmid containing optimal gRNA for each candidate GSH and the respective pMIA4.721 HDR-donor plasmids were nucleofected into hESC using the P3 Primary Cell kit (V4XP-3024) and programme CA-137. 1.5x10^6 cells were used for each targeting and were plated onto geltrex coated wells on 6-well plates in mTeSR with CloneR (Stemcell Technologies, 05889) following nucleofection. After 24h media was changed to mTeSR, and cells were allowed to recover for another 24-48h. Once cells reached 70-80% confluency, Blasticidin (ThermoFisher Scientific, A1113903) was added to the culture media at 10 µg/ml. Individual colonies were manually picked from the wells after 7-14 days of selection and expanded further for screening.

**Junction PCR**

Genomic DNA samples for all the collected GSH clones was isolated using PureLink™ Genomic DNA Mini Kit (ThermoFisher Scientific, K182002) according to manufacturer’s instructions. PCR reactions amplifying both 5’ and 3’ targeting HDR junctions as well as the wild type allele were set up using primers listed in Supp. Table 2. Samples were checked for the correct amplification size and alignment of the Sanger sequencing reads for each junction PCR and wild type allele.

**Off-target analysis**

The top five predicted off-targets were checked via PCR amplification and Sanger sequencing. PCR primers for respective off-targets for each gRNA are listed in Supp. Table 2. Sanger sequencing traces covering the off-target site for wildtype and the respective targeted clone are shown in Supp. Fig. 3.

**Copy number analysis**

We evaluated Blasticidin and RPP30 Copy Numbers using Droplet Digital™ Polymerase Chain Reaction (ddPCR) technology (Bio-Rad Technologies), according to manufacturer’s specifications. Briefly and following fluorescence-based quantification (ThermoFisher Scientific, Qubit™), 2.5 ng double-stranded DNA was added to a reaction mix containing target-specific primers/probe mixes (900 nM primer/250 nM probe per FAM and HEX fluorophore; Bio-Rad, 10042958 Unique Assay ID: dCNS626289650 and 10031243 Unique Assay ID: dHsaCP2500350), 0.05 U HaeIII Restriction Enzyme (New England Biolabs, R0108S) and ddPCR-specific Supermix for Probes (no dUTP) (Bio-Rad, 1863024). This was randomly partitioned into at least 10,000 discrete oil droplets per reaction using microfluidics within the QX200™ Droplet Generator (Bio-Rad, 1864002; together with Droplet Generation Oil for Probes, 1863005), which were gently transferred using a multi-channel pipette into a semi-skirted 96-well plate before heat-sealing (Bio-Rad PX1™ PCR Heat Sealer, 1814000). Target amplification within each droplet was conducted in the C1000 Touch™ Thermal Cycler with 96-Deep Well Reaction Module (Bio-Rad, 1851197) through the following PCR protocol: 1) Enzyme Activation at 95°C for 10 minutes, 2) 40 cycles of Denaturation and annealing/extension at 94°C for 30 seconds and 55°C for 1 minute, respectively, 3) Enzyme Deactivation at 98°C for 10 minutes. The QX200 Droplet Reader (Bio-Rad, 1864003) then derived the number of target-containing droplets through assessing each droplet for elevated, target-specific fluorescence. Blasticidin (FAM)-positive droplet counts were normalised using its respective well-specific RPP30 (HEX)-positive counts prior to downstream analysis. All experiments were done in duplicates, with data visualised and assessed using the QuantaSoft software version 1.7.4.917 (Bio-Rad).

**qPCR analysis**

RNA was extracted from three biological replicates of the GSH targeted H1 hESC Pansio-1, Olônne-18 and Keppel-19, control targeted cells (cells that underwent CRISPR/Cas9-mediated targeting and HDR of an expression cassette at a non-GSH locus) and two independent cultures of untargeted cells using Direct-zol^Tm^ RNA Miniprep kit (Zymo Research, ZYR.R2052). 1µg of RNA was converted into cDNA with Superscript IV Vilo MM (ThermoFisher Scientific, 11766050). Quantitative PCR reactions using TaqMan gene expression assays and master mix were used to compare the expression levels of *MAGI3*, *TXNL1* and *ZNRF4* against reference genes *18S* and *GAPDH* (ThermoFisher Scientific, Hs00326365_m1; Hs00169455_m1; Hs00741333_s1; HS99999901_S1; Hs03929097_g1 and 4444557). Quantitative RT-PCR analysis was done as described previously (Taylor et al., 2019) and resulting log2 fold change gene expression data was compared against reference H1 untargeted samples.

**RNA-seq library prep**

RNA samples described above for qPCR were also used for RNA-seq. RNA concentration and quality were checked with an Agilent 2100 RNA Pico Chip (Agilent, 5067-1513). RNA sequencing libraries were prepared using the TruSeq Stranded Total RNA Sample Prep Kit (Illumina, 20020596) including Ribo-Zero to remove abundant cytoplasmic rRNA. The remaining intact RNA was fragmented, followed by first- and second-strand cDNA synthesis using random hexamer primers. “End-repaired” fragments were ligated with a unique illumina adapters. All samples were multiplexed and pooled into a single library. Sequencing was done on a HiSeq 4000 to a minimum depth of 50 million 150 bp paired-end reads per biological sample.

**RNA-seq quality Control**

In all experiments, the raw paired-end reads in fastq format were initially processed with FastQC (<https://www.bioinformatics.babraham.ac.uk/projects/fastqc/>) for quality control at the base and sequence level. To remove the PCR duplicates we utilised the FastUniq algorithm (Xu et al., 2012). The adaptor trimming was performed by Trimmomatic (version 0.39) (Bolger et al., 2014). We quantified the 229,649 annotated human transcripts of GENCODE v35 by Kallisto (version 0.46) (Bray et al., 2016) followed by conversion of transcript to raw and TPM-normalized gene counts by the tximport package in R (Soneson et al., 2016). In total, 40,198 genes were quantified. Subsequently, we performed QC at the raw gene counts, checking for bad quality samples having less than 100,000 reads or more than 10% reads mapped to mitochondrial RNA or less than 2,000 detected genes. All samples of the various experiments were of high quality and were retained for the main analysis.

**RNA-seq Differential Expression Analysis**

The differential expression analysis was conducted by DEseq2 (Love et al., 2014) evaluating all pairwise comparisons of the conditions of each experiment. In each comparison, we considered only the expressed genes, i.e. those with non-zero raw counts in at least one sample. The differentially expressed genes were those with |logFC| ≥ 1 and FDR ≤ 0.01.

**Functional Enrichment Analysis**

Functional enrichment analysis was performed on the differentially expressed genes using the g:GOSt R package for g:Profiler (version e104_eg51_p15_3922dba) with g:SCS multiple testing correction method applying significance threshold of 0.05 (Raudvere et al., 2019).

**Karyotyping**

For each cell line, 20 GTL-banded metaphases were counted, of which a minimum of four have been analysed and karyotyped.

**Flow cytometry**

H1 wild type hESCs and Pansio-1, Olônne-18, and Keppel-19 H1 lines carrying Clover-transgene were disassociated with accutase (Stemcell Technologies, 07922) and resuspended in PBS. The single cells in PBS were analysed with a BD LSR Fortessa x-20 FACS Analyzer and FlowJo (v10.6.1).

**hESC cardiac differentiation**

Two days prior to starting differentiation, cells were dissociated using Accutase and seeded as single cells in Geltrex-coated 12-well plates at seeding density between 1 to 1.5x10^6 cells. Cardiac differentiation was performed following the published protocol by Lian et. al. (Lian et al., 2013), with modifications as follows. 6 µM of CHIR99021 (Stemcell Technologies, 72054) was added on day 0 and left for 24h followed by medium change. On day 3, 5µM IWP2 (Sigma Aldrich, I0536) was added using 50/50 mix of new fresh medium and conditioned medium collected from each well and left for 48h. Culture medium from day 0 until day 7 was RPMI1640 (HyClone, SH30027.01) plus B-27 serum-free supplement without insulin (Gibco, A1895601). From day 7 and onwards RPMI1640 with B-27 serum free supplement with insulin (Gibco, 17504044) was used and changed every 2-3 days.

**hESC differentiation to hepatocyte-like cells**

hESCs were differentiated into hepatocyte-like cells as described previously (Hannan et al., 2013; Ng et al., 2019), with some modifications. Briefly, hESCs were dissociated into small clumps using ReLeSR and plated onto gelatin-coated coverslips in a 12-well plate with mTeSR. Two days later, hESCs were induced to differentiate into definitive endoderm (DE) cells in RPMI-1640 medium (Gibco) containing 2% B-27 (Invitrogen), 1% non-essential amino acids (Gibco), 1% GlutaMAX™ (Gibco) and 50μM 2-mercaptoethanol (Gibco) (basal differentiation medium), supplemented with 100ng/ml Activin A (R&D Systems), 3 μM CHIR99021 (Tocris) and 10 μM LY294002 (LC Labs) for the first three days (D0 to D3). From D3 to D6, cells were incubated in basal differentiation medium supplemented with 50 ng/ml Activin A to form foregut endoderm cells. From D6 to D10, cells were incubated in basal differentiation medium supplemented with 20 ng/ml BMP4 (Miltenyi Biotec) and 10ng/ml FGF10 (Miltenyi Biotec) to form hepatic endoderm cells. From D10 to D24, hepatic endoderm cells were incubated in HCM Bulletkit (Lonza) differentiation media supplemented with 30 ng/ml Oncostatin M (Miltenyi Biotec) and 50 ng/ml HGF (Miltenyi Biotec). Differentiation medium was replaced every two or three days.

**hESC neural induction**

H1 cells cultured in mTeSR complete medium for 1-2 days were then used for neural induction as published (Li et al., 2011; Wang et al., 2017). Briefly, 20-30% confluent H1 cells were treated with CHIR99021, SB431542 and Compound E in neural induction media, changed every 2 days; 7 days later, the cells were split 1:3 by Accutase and seeded on matrigel-coated plates. ROCK inhibitor (1254, Tocris) was added (final concentration 10μM) to the suspension at passaging. Cells were then cultured in neural cell culture medium. These derived cells are neural precursor cells (NPC), which were used for further studies.

**Neuronal differentiation**

Spontaneous neuronal differentiation was performed as previously described (Li et al., 2011). Briefly, the derived 2 X10^5^ NPCs were seeded on poly-l-lysine (P4707, Sigma) and laminin (L2020, Sigma) coated 6-well plates in neural cell culture medium. The next day, the cells were cultured in neuron differentiation medium: DMEM/F12 (11330-032), Neurobasal (21103-049), 1 X N2 (17502-048), 1 X B27 (17504-044), 300ng/ml cAMP (A9501), 0.2mM vitamin C (A4544-25), 10ng/ml BDNF (450-02), 10ng/ml GDNF (450-10) until day 30.

**Immunofluorescence**

Cells on coverslips were fixed in 4% paraformaldehyde (Wako) for 15 min at room temperature, before blocking in 5% donkey serum (EMD milipore) in PBS with 0.1% Triton X-100 for 1h at room temperature. Cells were stained with primary antibody overnight at 4°C (see Supp. Table 2 for antibodies used), or for control slides with blocking buffer. Secondary antibody staining was done with the appropriate AlexaFluor 594 for 1h at room temperature. Lastly, cells were stained with DAPI (Sigma-Aldrich, 1:5000) for 20 min at room temperature. Coverslips were mounted onto glass slides using Vectashield (Vector Laboratories). Images were taken using the EVOS M5000 microscope. Light intensity and gain were kept consistent across samples and controls with each antibody.

**Author contributions**

Conceptualisation, M.I.A.; Methodology, M.I.A.; Software, M.I.A., E.M., and T.B.A.; Formal analysis, M.I.A.; E.M., and R.S.Y.F; Investigation, M.I.A., A.P., V.D.D., J.W., J.T., Z.T. and W.X.T.; Resources, J.W. and A.K.K.T; Writing – Original Draft, M.I.A. and R.S.Y.F; Writing – Review & Editing; all authors; Supervision, M.I.A., A.K.K.T and R.S.Y.F; Funding Acquisition, M.I.A.

**Acknowledgements**

This manuscript is dedicated to the memory of Seppo J. Autio. We would like to express our gratitude to Dr Alexander Lezhava for generously supporting our ddPCR experiments. We thank the late Albert Barillé for life-long inspiration. We would also like to thank Dr Steve Oh for insightful discussions and feedback on the manuscript, as well as Dr Lee Siggens for feedback and assistance on primer design. This research was funded by the BMRC YIG 2016 (1610851033) and A*STAR CDF 2020 (202D8020) to M.I.A.. The funding bodies played no role in the design of the study and collection, analysis, and interpretation of data and in writing the manuscript.

**Declaration of interests**

The authors declare no competing interests.

**Supplementary Figure 1.** Screenshots of Hi-C interaction matrices from H1 hESC for each of the shortlisted GSH candidate loci. TADs are indicated by the “pyramids” of high interaction observed in the Hi-C matrices. UCSC genome browser track annotating the candidate GSH (targeted GSH in pink), GENCODE v36 and H3K27Ac mark from ENCODE shown below.

**Supplementary Figure 2.** PCR gel images of junction PCR and wild type allele PCR reactions for screened clones.

**Supplementary Figure 3.** Sanger sequencing traces of the top five predicted gRNA off-target sites for un-targeted wild type H1 hESC and the respective GSH targeted clones.

**Supplementary Figure 4.** Representative images of Pansio-1 line immunofluorescence staining with AF594 secondary antibody alone. DAPI = nuclear staining with 4′,6-diamidino-2-phenylindole, Clover = fluorescence from Clover transgene, AF594 = staining with secondary antibody alone, Overlay = overlay of the three imaged channels.
