## Supplementary material for "Computationally defined and *in vitro* validated putative genomic safe harbour loci for transgene expression in human cells": supp fig 1

#### Targeted GSH on chr1

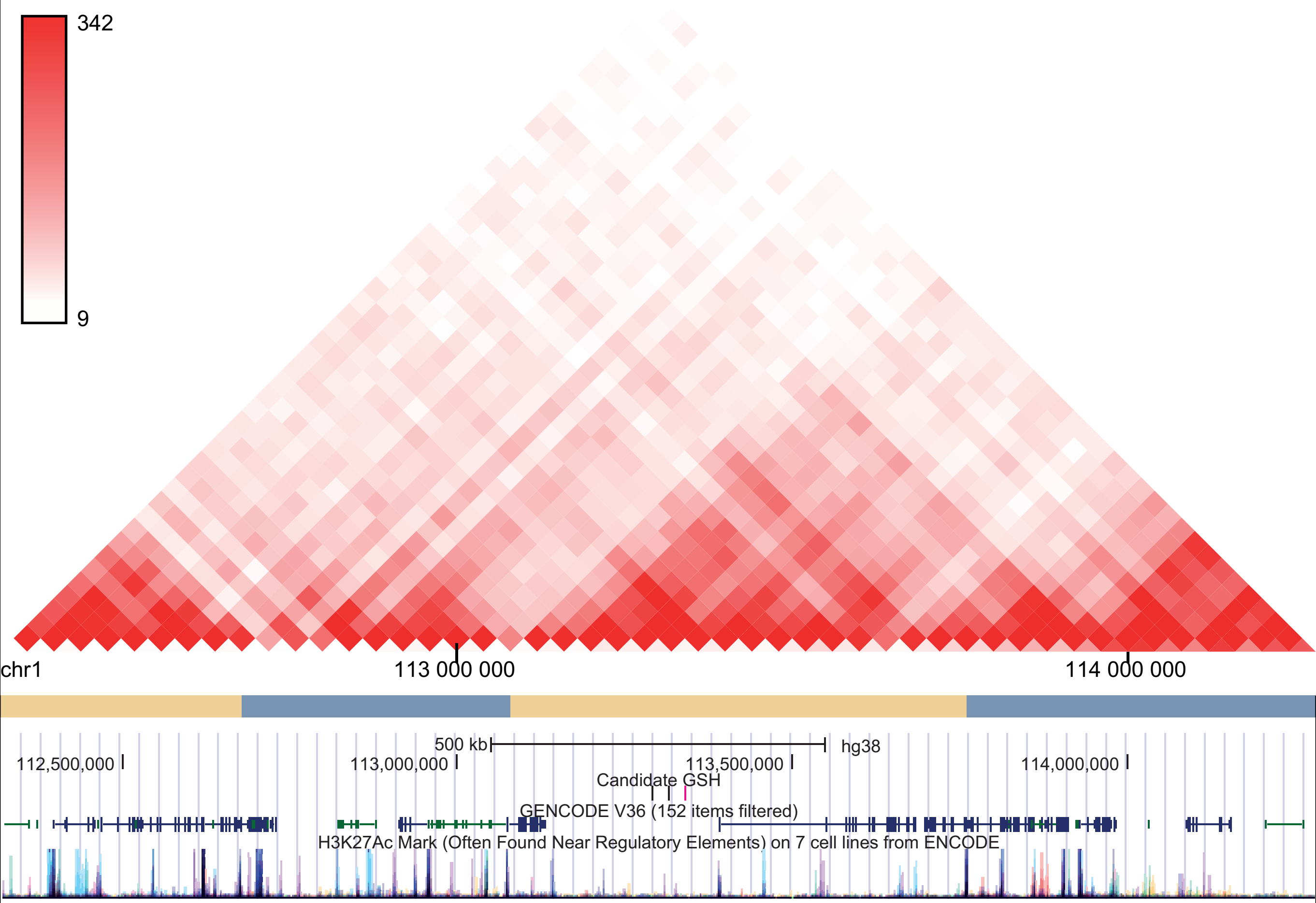

### Targeted GSH on chr2

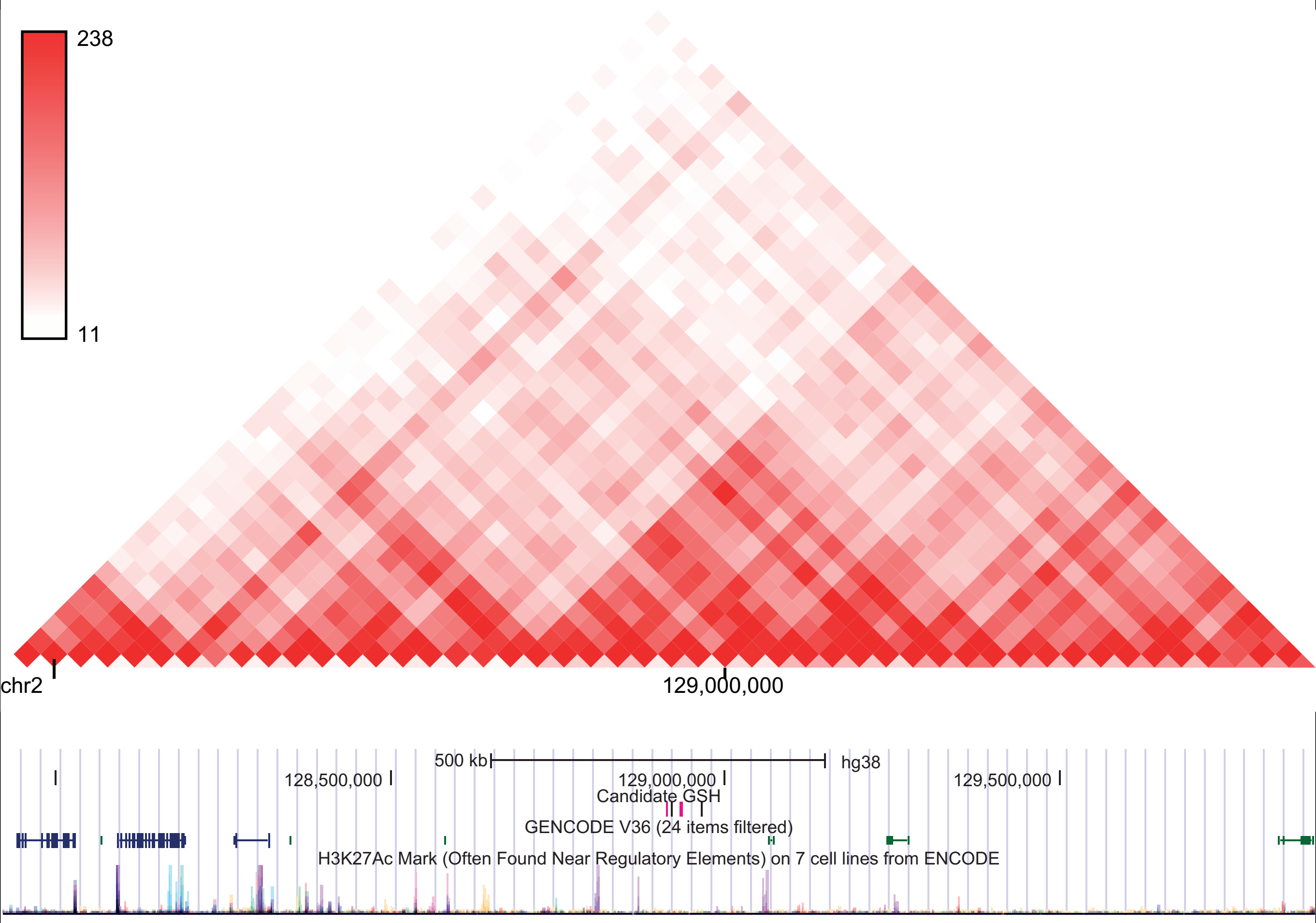

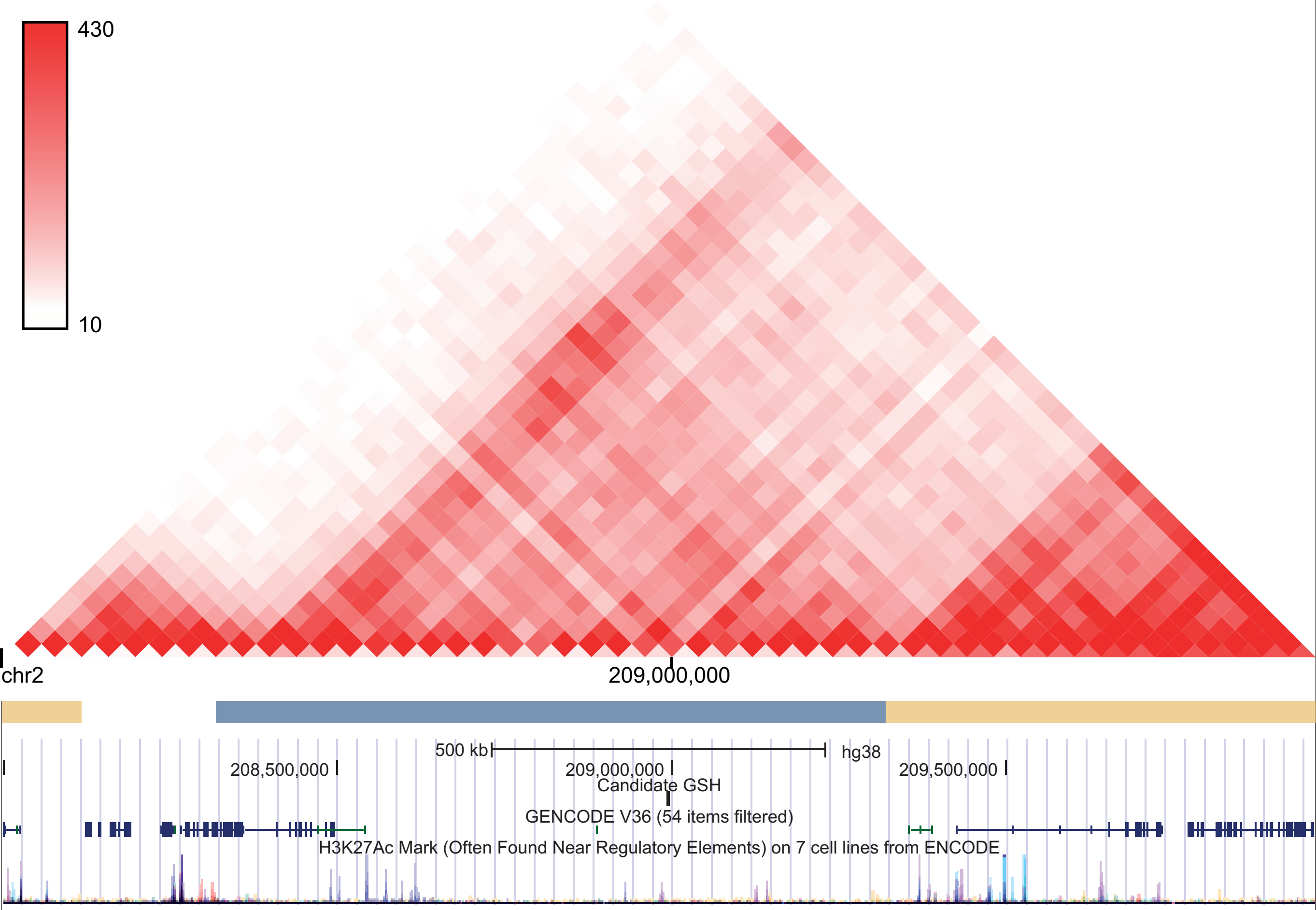
430
10
chr2
209,000,000
500 kb
hg38
208,500,000
209,000,000
209,500,000
Candidate GSH
GENCODE V36 (54 items filtered)
H3K27Ac Mark (Often Found Near Regulatory Elements) on 7 cell lines from ENCODE

### Targeted GSH on chr4

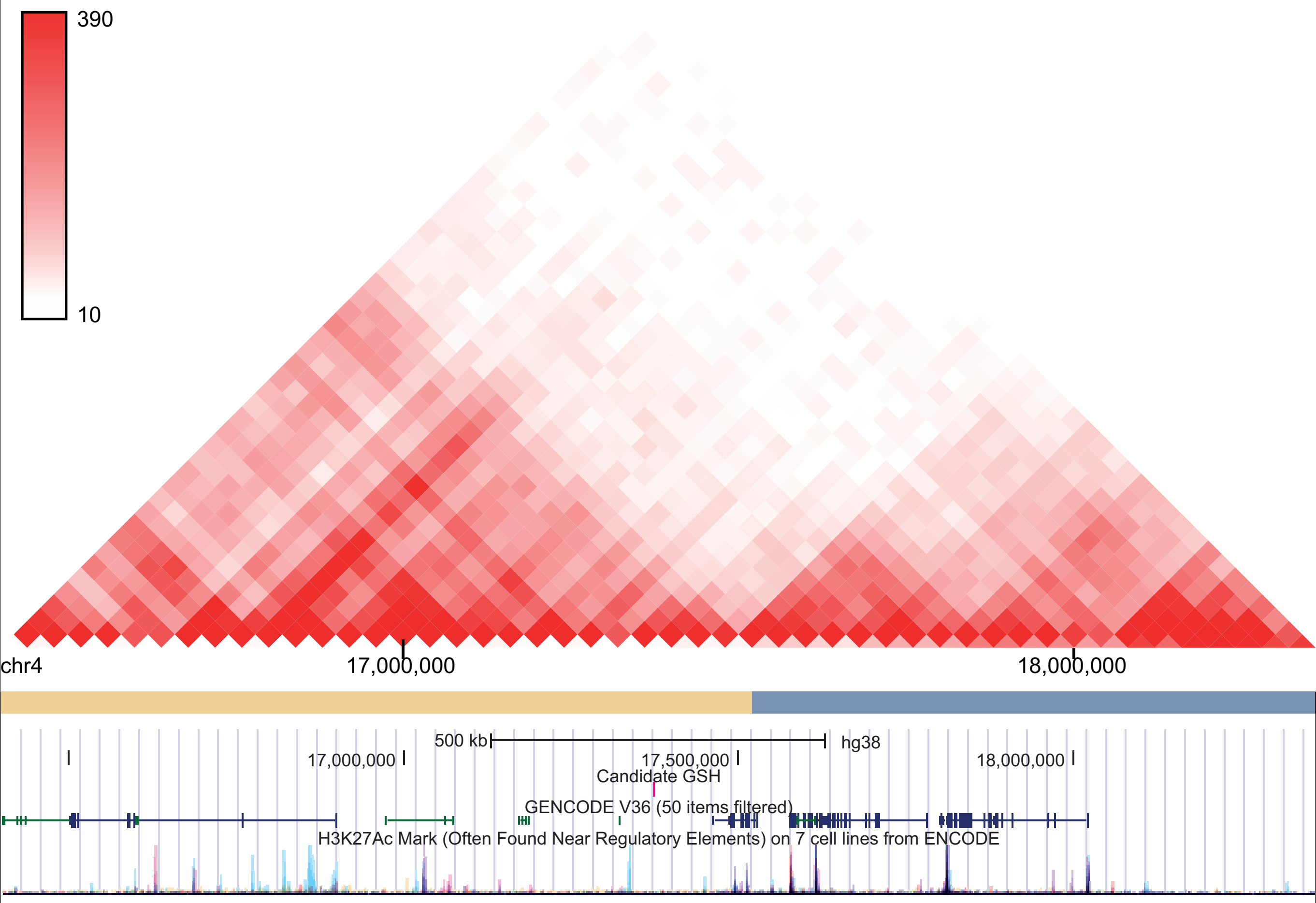

### Targeted GSH on chr6

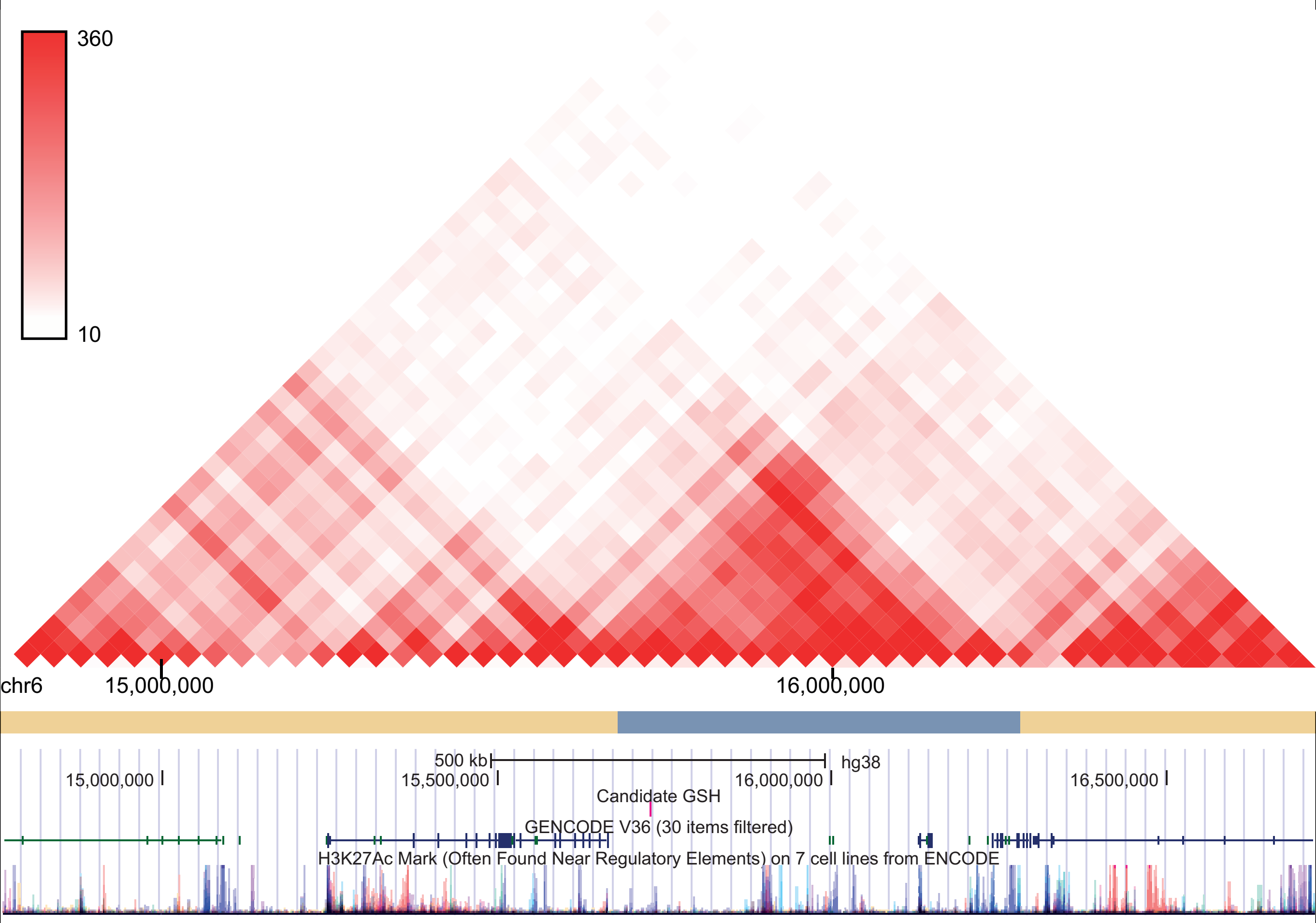

### Targeted GSH on chr18

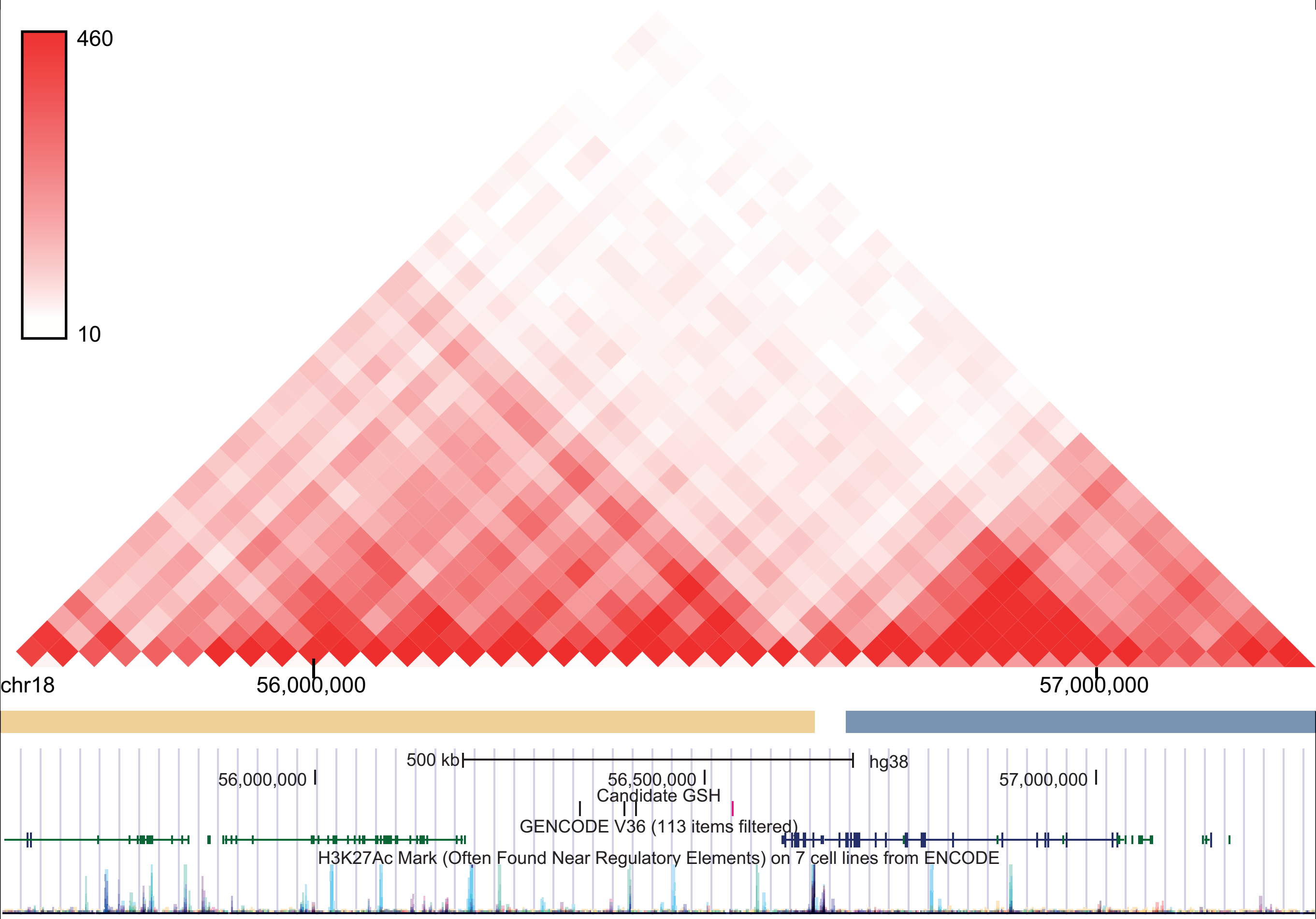

### Targeted GSH on chr19

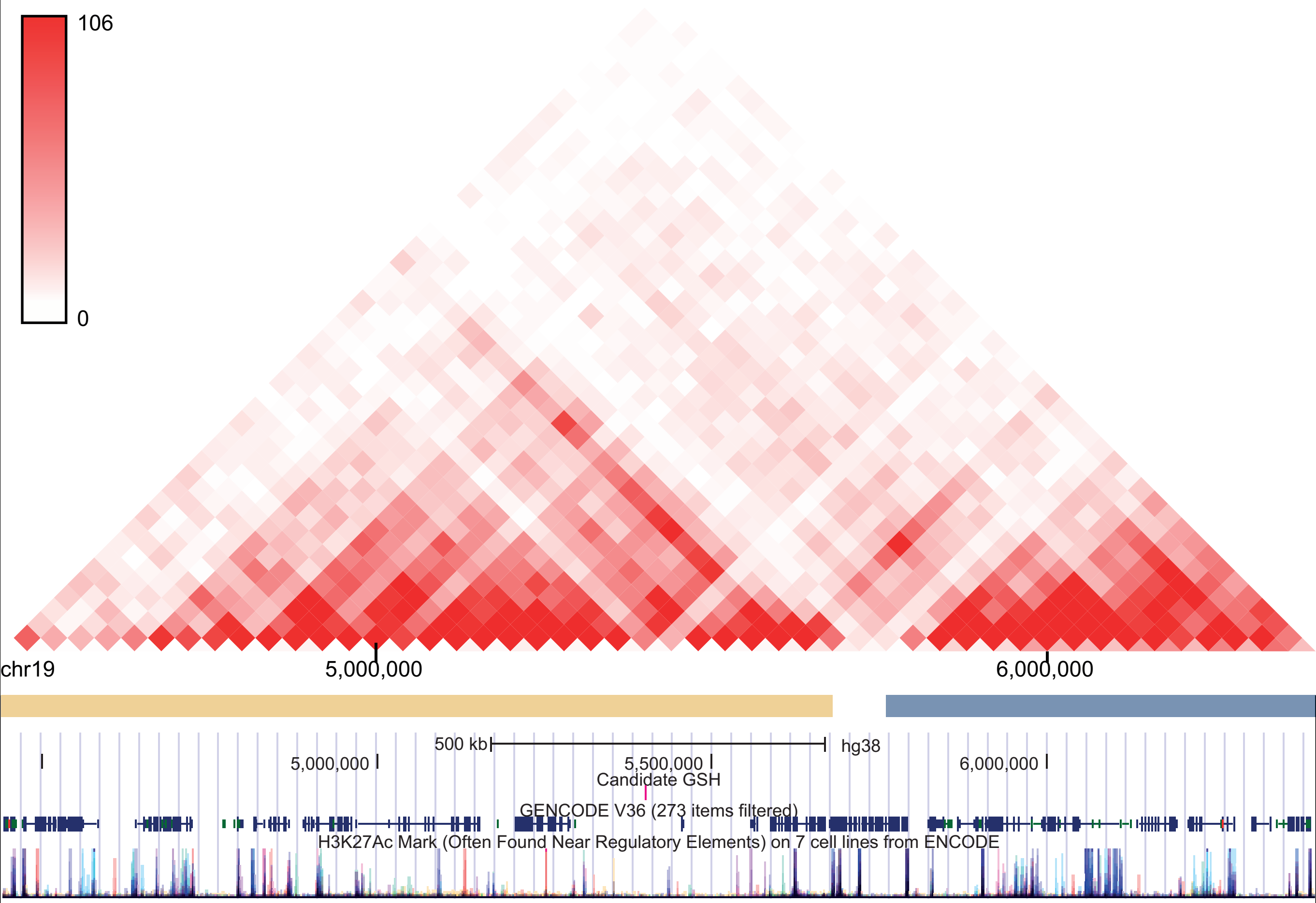
