## Supplementary figures and images for "Computationally defined and *in vitro* validated putative genomic safe harbour loci for transgene expression in human cells"

### supp fi 3

A

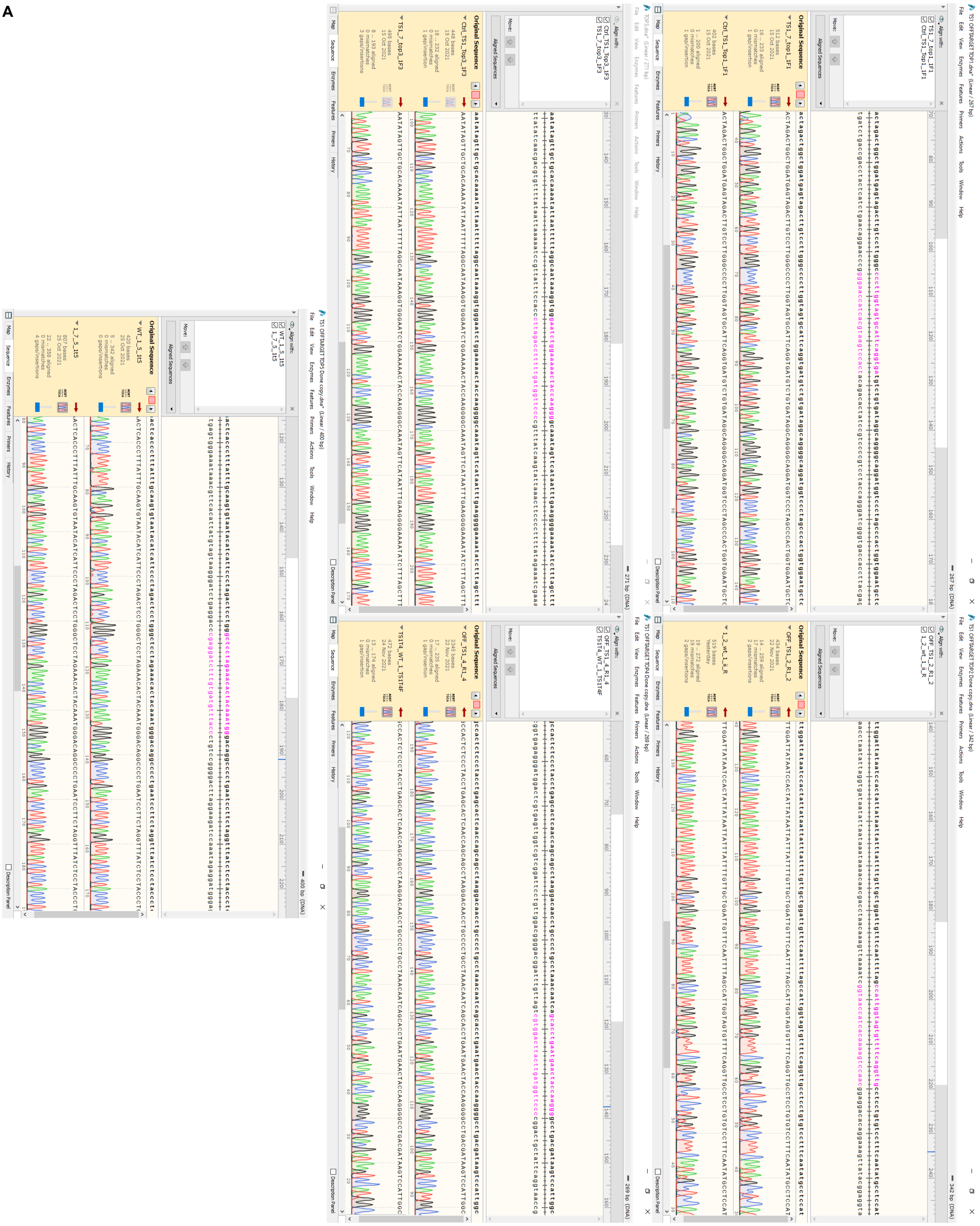

B

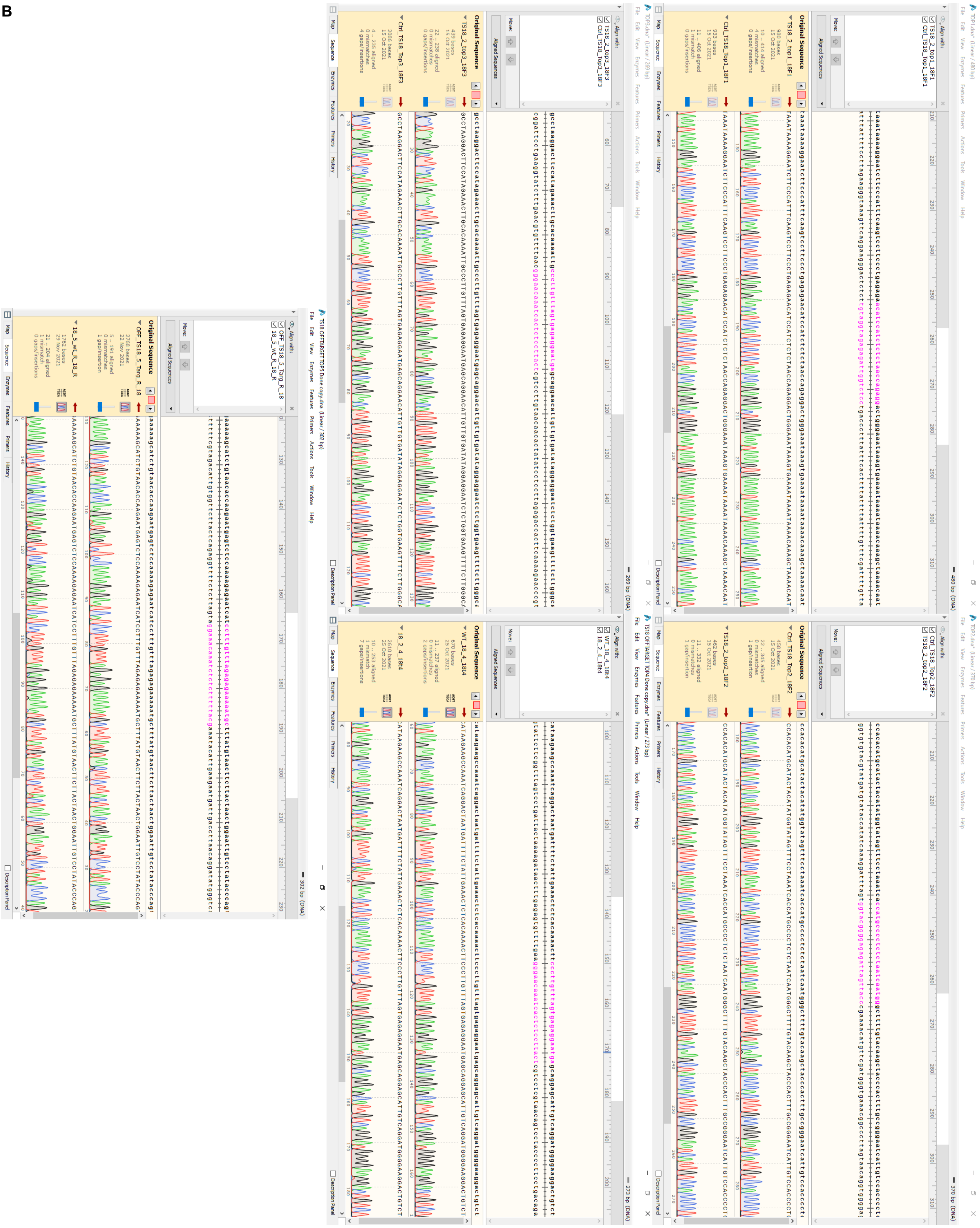

C

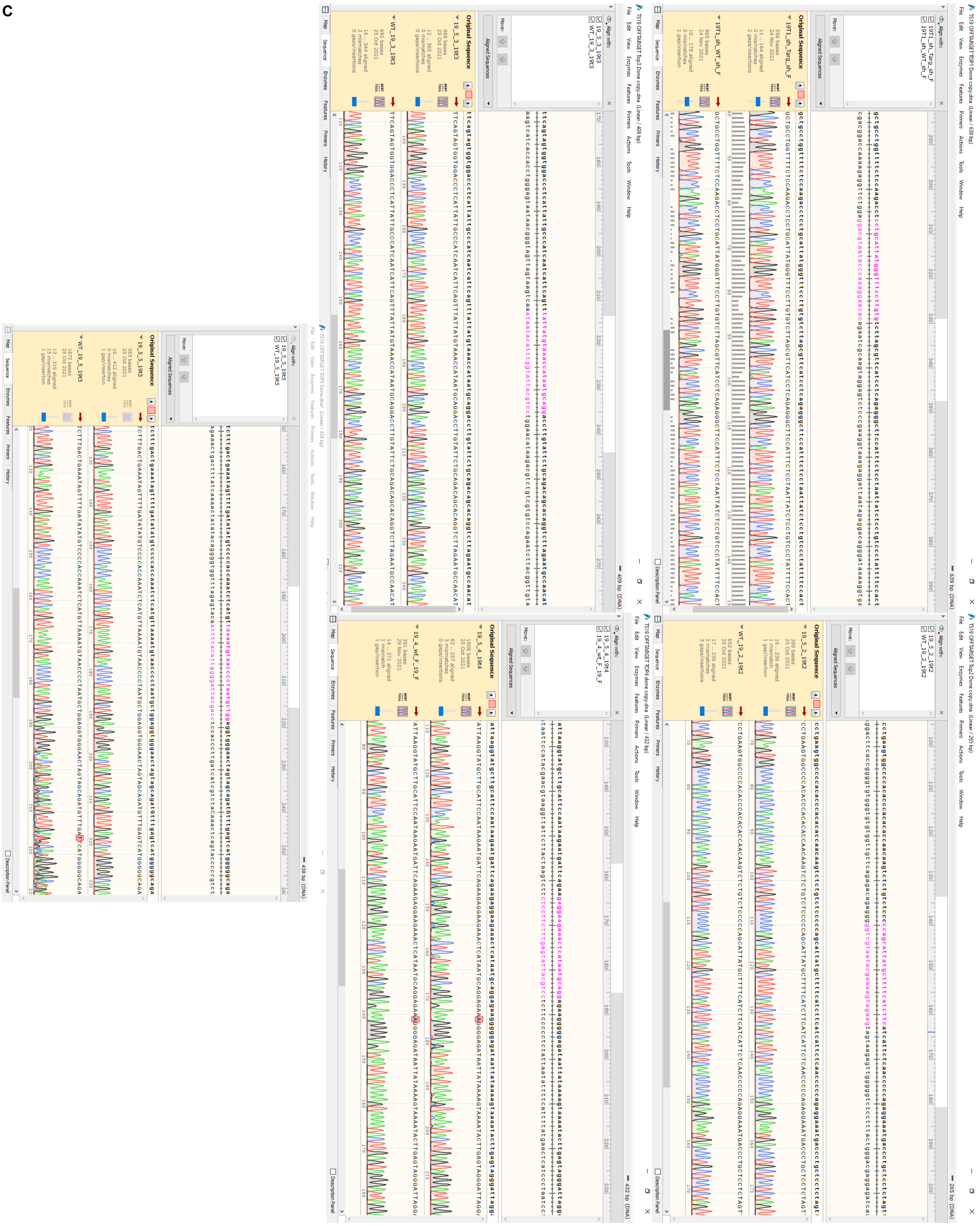

### supp fig 2

#### Pansio-1

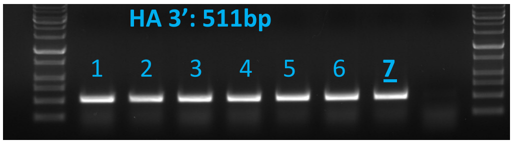


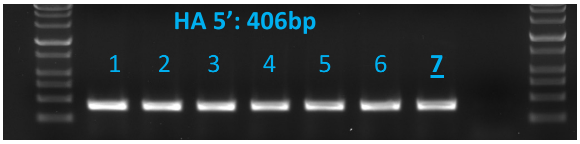


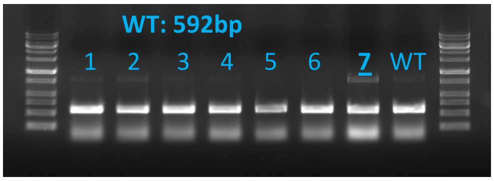


#### Olônne-18

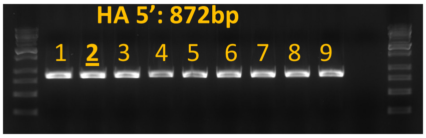


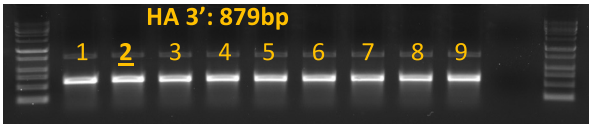


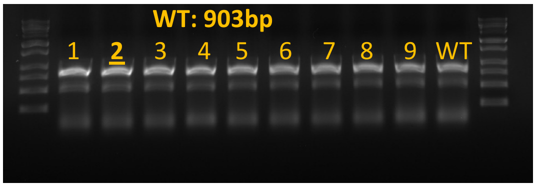


#### Keppel-19

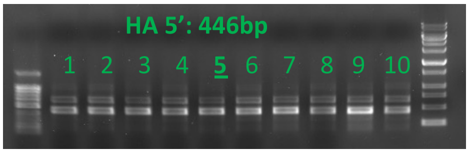


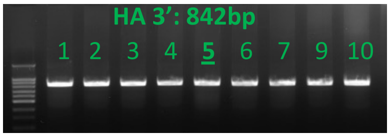


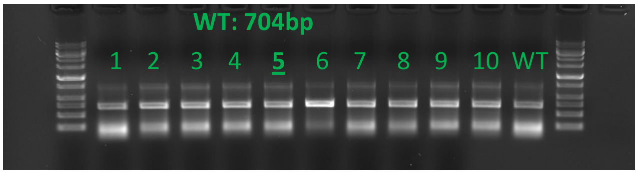

### supp fig 4

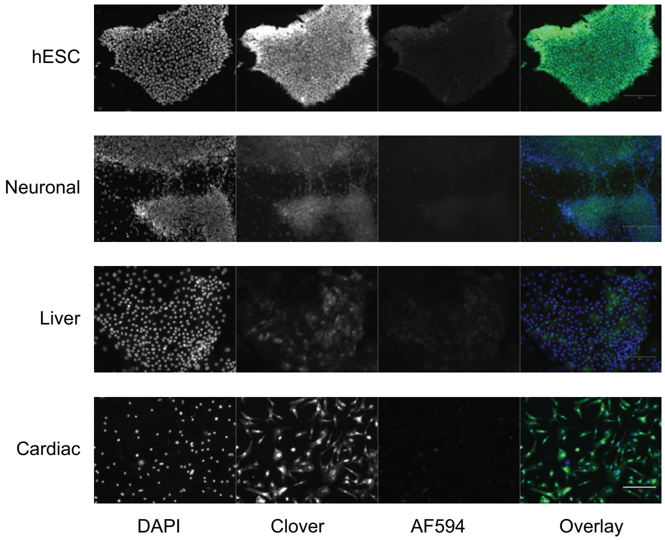
